# Humans forage for reward in reinforcement learning tasks

**DOI:** 10.1101/2024.07.08.602539

**Authors:** Meriam Zid, Veldon-James Laurie, Jorge Ramírez-Ruiz, Alix Lavigne-Champagne, Akram Shourkeshti, Dameon Harrell, Alexander B. Herman, R. Becket Ebitz

**Author notes:** These authors contributed equally to this work (Alexander B. Herman, R. Becket Ebitz).

## Abstract

How do we make good decisions in uncertain environments? In psychology and neuroscience, the classic view is that we calculate the value of each option, compare them, and choose the most rewarding modulo exploratory noise. An ethologist, conversely, would argue that we commit to one option until its value drops below a threshold and then explore alternatives. Because the fields use incompatible methods, it remains unclear which view better describes human decision-making. Here, we found that humans use compare-to-threshold computations in classic compare-alternative tasks. Because compare-alternative computations are central to the reinforcement-learning (RL) models typically used in the cognitive and brain sciences, we developed a novel compare-to-threshold model (“foraging”). Compared to previous RL models, the foraging model better fit participant behavior, better predicted the tendency to repeat choices, and predicted held-out participants that were almost impossible under compare-alternative models. These results suggest that humans use compare-to-threshold computations in sequential decision-making.

## Introduction

Because the world is only imperfectly observable and changes frequently, many of the decisions we make are uncertain. How do we make good decisions in the presence of this uncertainty? The conventional view in the cognitive and brain sciences is that we (1) estimate the subjective utility or “value” of each option being considered and then (2) compare alternative option values to make the best possible decision [1–5]. This “compare-alternatives” perspective is ubiquitous in psychology and neuroscience. For example, nearly all **Reinforcement Learning** (RL) models, from the earliest Rescorla-Wagner model [1] to more modern models like Q-Learning and SARSA [6, 7], assume that the value of choosing each alternative option are compared in some way in order to select the best action [1, 4, 6]. These models, as well as the compare-alternatives perspective more broadly, shape the way that tasks are designed in these fields. For example, because we assume that values must be compared against each other, laboratory tasks using cued rewards often present all options simultaneously [8–11]. Similarly, because the reward prediction error in RL models functions to track changing mean values, reward contingencies in reward learning tasks tend to both increase and decrease over time [12–15]—in contrast decay of exploited resources often seen in natural environments.

While the compare-alternatives view is widely influential, it is not universally accepted. In ethology, for example, **foraging** models instead assume that we (1) calculate the value of one exploited option and (2) compare this one value against a threshold to decide whether to continue exploiting or try something new [16, 17]. The foraging view grew out of natural environments, where the value of an exploited resource will tend to decay over time and options are encountered sequentially, rather than in parallel. As a result, although foraging models have begun seeing some use among human cognitive scientists [18–23], these studies develop tasks that mirror the assumptions at the heart of the foraging perspective: they introduce reward contingencies that decay at a predictable rate (rath(er than changing unpredictably) and/or options that are encountered in sequence (rather than being presented simultaneously) [18–23]. In short, the compare-alternatives and compare-to-threshold views of sequential decision-making use incomparable formalisms to make sense of behavior in incompatible environments. As a result, we do not know which view best describes human decision-making in standard laboratory tasks.

Here, we asked whether human decision-making was better described as a compare-to-threshold process or as a compare-alternatives process in a classic testbed of decision-making under uncertainty from the reinforcement learning (RL) literature: a restless k-armed bandit task. We found that human behavior more closely resembled compare-to-threshold computations than compare-alternatives computations. This insight is difficult to reconcile with traditional RL algorithms from psychology and neuroscience, which model action selection with the critical assumption of compare-alternatives computations. Therefore, we developed a novel compare-to-threshold sequential decision-making model that is inspired by foraging theory rather than the compare-alternatives approach at the heart of RL. Across 3 independent experiments, we compared this new foraging model with various variations on traditional compare-alternatives RL models [4]. We found that the foraging model was a better fit for the participants’ decisions, outperformed multiple variations on traditional RL models, and better predicted the participants’ tendency to repeat choices on both individual and group level. Together, these findings suggest that humans use compare-to-threshold computations—even in tasks that are commonly used as testbeds for compare-alternatives computations.

## Results

### Participants performed above chance and used complex strategies

In Experiment 1, participants on the Amazon mTurk platform (258 people, 120 female, 2 other or non-reporting) performed a classic sequential decision-making task known as a restless k-armed bandit (Figure 1A; [6, 12, 14, 15, 24, 25]). The participants were asked to repeatedly choose between k options, each of which was associated with some probability of reward. Reward probabilities changed unpredictably over time and independently across options (Figure 1B). The reward structure was not cued to the participants so the only way to infer the value of an option was to sample it. Because rewards evolved over time, the longer the participants went without sampling an option, the more uncertain its payoff became. This task naturally encourages decision-makers to alternate between exploiting valuable options and exploring uncertain alternatives because the latter could become better at any time. It is a classic in the RL literature and has become a testbed for evaluating RL models in both psychology and artificial intelligence (AI) [14, 26, 27].

**Fig. 1:**
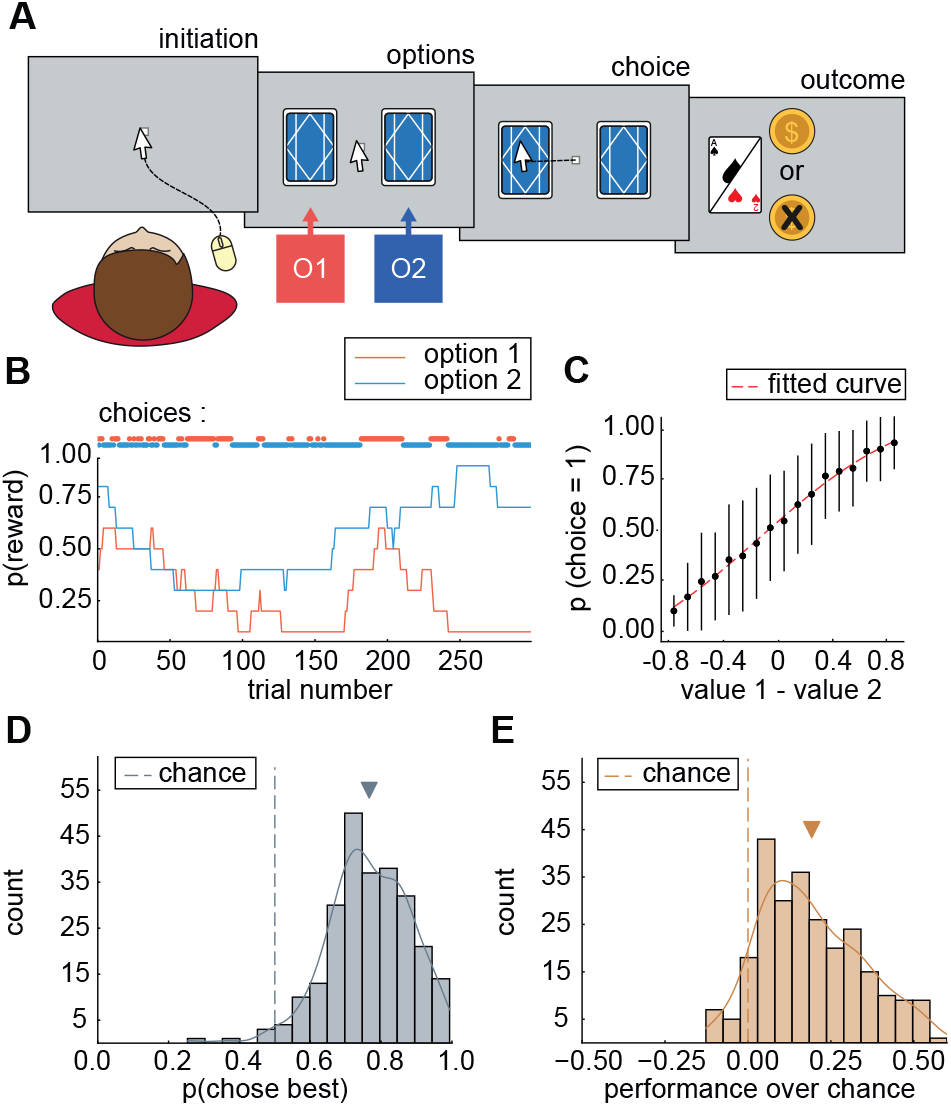
Task and baseline behavior. **A)** Participants (n=258) chose between 2 face-down decks of playing cards that were composed of aces and 2’s. Participants were told that the proportion of aces and 2’s in each deck would change over time and that decks could be good, bad, or mediocre right now, but would not stay that way the whole time. Participants got a point (and $0.02) for each ace they found, but no points for 2’s. **B)** The reward probability of each option (red trace = option 1; blue trace = option from an example reward schedule, with participant choices along the top (red dots = option 1; blue dots = option 2). **C)** Average probability of choosing option 1 as a function of the difference in the objective reward probability (“value”) between option 1 and 2. Error bars = standard error of the mean (SEM) across participants. **D)** Distribution of the number of trials in which each participant chose the objectively best option. Dotted line = chance, caret = mean across participants.**E)** Same as *D*, for the percent of trials in which participants were rewarded, normalized such that units represent percent of chance performance (caret = mean).

Participants were generally good at the task (Figure 1C), despite its uncertainty. They chose the objectively best option 76.6% of the time (+/-11.5% STD; Figure 1D) and received rewards 19.2 % more frequently than would be expected by chance (+/-15.3 % STD; Figure 1E). (Note that 4/258 participants [1.6%] were excluded from this and further analyses because they chose only one option.) Participants were more likely to repeat choices than to switch (switching on only 19.9% of trials, +/-14.5% STD). They were also more likely to repeat after reward (win-stay = 93.3%, +/-11.1% STD) and switch after no-reward (lose-shift = 39.2%, +/-21.0% STD). However, no participants followed a deterministic win-stay-lose-shift rule (all participants either win-shifted or lose-stayed at least 5% of the time).

### Human behavior better resembles the fingerprint of compare-to-threshold decisions

Although both compare-alternatives and compare-to-threshold views of sequential decision-making come from different fields, they describe the same behavior: exploiting high-value options when those are available, but also occasionally switching to alternatives that could be better. Where the compare-alternatives and compare-to-threshold views differ is in how the decision to switch between options is made. As a result of these differences, compare-alternatives and compare-to-threshold views predict that switching will be most frequent in different kinds of environments (Figure 2A-C).

**Fig. 2:**
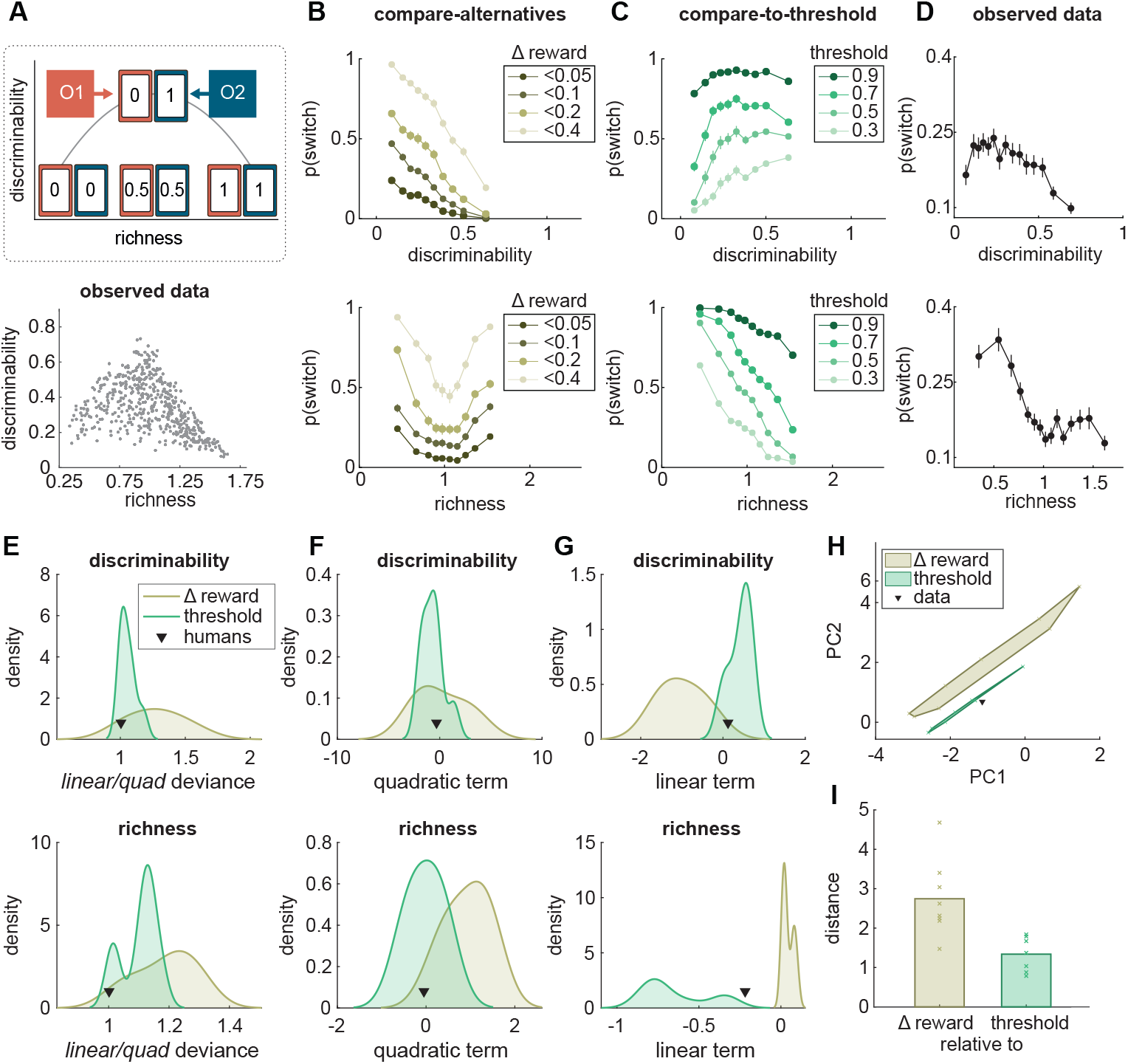
Compare-to-threshold and compare-alternatives decisions have dissociable signatures in rich and discriminable environments. **A)** *top*: Cartoon illustrating the non-linear relationship between reward discriminability (*i*.*e*., how different the reward probability of all options are) and richness (*i*.*e*., How rewarding all the options are) in a bounded environment. *bottom*: discriminability as a function of richness in Experiment 1. Each dot represents a segment of 100 trials experienced by one participant. **B)** In compare-alternatives decisions, switching behavior is maximal when the absolute difference in reward probability (Δ reward) is low [24]. If we take the difference in rewards as a proxy for the probability of switch, then the compare-alternatives hypothesis makes different predictions for how switching should change as a function of reward discriminability (*top*) and richness (*bottom*). Shades of gold illustrate different hypothetical “thresholds” for switching meaning that if the difference in reward probabilities between all options (Δ reward) is below these hypothetical threshold the decision is to switch options. **C)** In compare-to-threshold decisions, switching behavior is maximal when the sum of rewards is low. Here, each option’s value is individually compared to a preset threshold. If the average of these below-threshold comparison serves as a proxy for switching then the compare-to-threshold hypothesis makes different predictions for how switching should change as a function of reward discriminability (*top*) and richness (*bottom*). Shades of green illustrate different hypothetical “thresholds” for switching meaning that if the option’s reward probability is below these hypothetical threshold the decision is to switch options. **D)** Probability of switch of all participants in Experiment 1 as a function of different levels of reward discriminability *(top)* and richness *(bottom)* of the environment (errorbar = SEM). **E)** Distribution of deviance ratio between linear and quadratic model fits to compare-alternatives (*gold*) and compare-to-threshold (*green*) proxies (Figure2B-C) as a function of reward discriminability *(top)* and richness *(bottom)* of the environment (black caret = humans). **F)** Distribution of the quadratic term (*i*.*e*., curvature) of the quadratic model fit to compare-alternatives (*gold*) and compare-to-threshold (*green*) proxies (Figure2B-C) as a function of reward discriminability *(top)* and richness *(bottom)* of the environment (black caret = humans). **G)** Distribution of the linear term (*i*.*e*., slope) of the linear model fit to compare-alternatives (*gold*) and compare-to-threshold (*green*) proxies (Figure2B-C) as a function of reward discriminability *(top)* and richness *(bottom)* (black caret = humans). **H)** Principle components (PC) projection of the multidimensional features of the compare-alternatives (*gold*) and compare-to-threshold (*green*) proxies (see Methods). Bounds encompass all simulations. x’s = individual simulations, black caret = humans. **I)** Distance between humans and every compare-alternatives (*gold*) and compare-to-threshold (*green*) proxy in the multidimensional space. Bars = mean distance, x’s = individual simulations.

The compare-alternatives view predicts that people should switch options more often when it is most difficult to discern which option is the best. Because switching occurs most frequently when option values are close together, people should switch more frequently in **ambiguous** environments versus *discriminable* environments (Figure 2B-*top*). The compare-to-threshold view, conversely, predicts that people should switch more whenever an exploited option falls below a threshold. Because the value of an option is more likely to fall below threshold when all option values are low, people should switch more frequently in **poor** environments (versus *rich* environments) (Figure 2C-*bottom*). (*N*.*B*. We assume that the threshold is learned over long time scales such that it is essentially fixed within any given experiment [17], rather than adapting at a rate that would cause it to start approximating comparealternatives [21].) In short, switching should depend on very different aspects of the environment in compare-to-threshold decision-making versus compare-alternatives decision-making.

If we could dissociate richness from discriminability, we could identify whether people were using compare-to-threshold or compare-alternative computations simply by looking at which one influences switching behavior. This is because richness would not affect switching in compare-alternatives decisions and discriminability would not affect switching in compare-to-threshold decisions. However, when rewards are probabilistic, these variables are not orthogonal (Figure 2A). The environment can only be very poor (reward probability of all options close to 0) or very rich (all reward probabilities close to 1) when discriminability is low. This means that compare-alternatives decision-making will have a relationship with richness in this task, but that relationship will be U-shaped with minimal switching at intermediate levels of richness (where discriminability is highest; Figure 2B,*bottom*). This also means that compare-to-threshold decision-making will have a nonlinear relationship with discriminability. This relationship would be subtly inverted U-shaped, with maximal switching at intermediate levels of discriminability (where richness is lowest; Figure 2C,*top*). Thus, although compare-to-threshold decisions are not based on discriminability and compare-alternatives decisions are not based on richness, the fact that rewards are bounded means that each strategy should have a unique fingerprint as a function of these two environmental variables (Figure 2 B-C).

To determine whether human decision-making more closely resembled the predictions of the compare-alternatives or compare-to-threshold hypotheses, we therefore asked whether humans resembled the compare-alternatives or compare-to-threshold fingerprints. We found that the participants switched options the most at intermediate, rather than low levels of discriminability (Figure 2D,*top*), consistent with compare-to-threshold computations. The participants were also less likely to switch in rich environments, compared to poor ones (Figure 2D,*bottom*), again consistent with compare-to-threshold computations. To quantify the relationship between participants’ switching behavior and the respective fingerprints of the compare-alternatives and compare-to-threshold views, we simulated data at every possible threshold for compare-to-threshold decision-making and at every possible Δ reward for compare-alternatives decision-making. We then used generalized linear models (GLMs) to characterize the slope and curvature of these relationships for each hypothesis (Figure 2E-I; see Methods). Human switching was sensitive to discriminability and richness in a way that better resembled compare-to-threshold decision making, both in terms of every individual feature (2E-G) and in a multivariate feature space that simultaneously considered all features (2H-I; average distance to compare-alternative proxies = 2.75 +/-0.97 STD; to compare-to-threshold proxies = 1.34 +/-0.44 STD; t(13) = 3.51, *p <* 0.005; see Methods). In sum, while the human data was not a perfect match for either fingerprint, it better resembled the fingerprint of compare-to-threshold decision-making than the fingerprint of compare-alternatives decision-making.

### Compare-alternatives agents outperformed compare-to-threshold agents on the task

Although the k-armed bandit is a classic testbed for compare-alternatives models like traditional RL models [4, 6, 14, 28], it was nonetheless possible that some aspect of the task design encouraged the participants to use compare-to-threshold computations here. To determine if this was the case, we next developed a sequential decision-making agent based on compare-to-threshold computations and compared its performance on the task against a traditional compare-alternatives RL agent (Rescorla-Wagner; [6, 7]).

A traditional RL agent estimates the value of choosing each individual option (*V*_1_, …, *V*_*n*_; Figure 3A) and compares values across the option set (*compare-alternatives*; Figure 3A) to select the best possible action. Our novel foraging agent updates the value of the selected action via the same delta-rule computations used in a traditional RL agent, but differs in what the action represents. Specifically, the foraging agent only estimates the value of staying with the currently exploited option (*V*_exploit_; Figure 2B). It then compares this value against a fixed threshold to decide whether to continue exploiting or else to explore the alternatives at random (*compare-to-threshold*; Figure 3B).

**Fig. 3:**
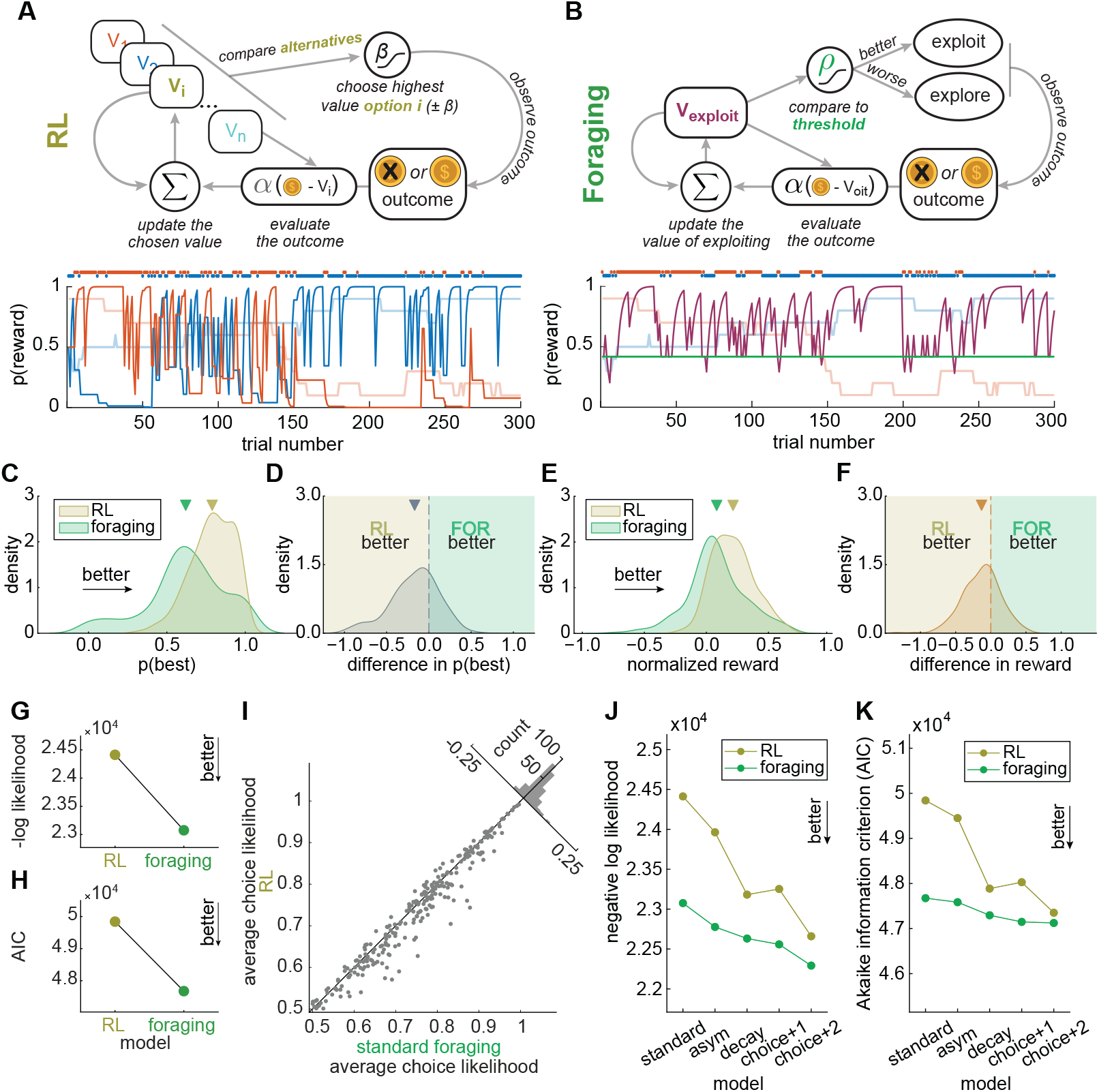
RL and foraging models produce different choice patterns performance and quality fits to the data. **A)** *Top*: A cartoon illustrating the mechanics of the traditional RL model. The model estimates the value of choosing each individual option through a standard delta-rule update. The value of choosing alternatives options are then compared to each other and the highest value is selected. *Bottom*: Simulated data from the traditional RL model, fit to an example participant from Experiment 1 and run on the corresponding reward schedule. The choices (*colored dots*) and estimated values (*colored traces*) of the traditional RL model are shown, along with the objective reward probabilities (*pale traces*). **B)** A cartoon illustrating the mechanics of the foraging model. The model estimates the value of staying with the currently exploited option through a standard delta-rule update. The value of exploitation is then compared to a threshold in order to decide whether to stay and continue exploiting or to switch and explore other options. *Bottom*: Simulated data from the foraging model, fit to an example participant from Experiment 1 and run on the corresponding reward schedule. The choices (*colored dots*) and exploitation value (*purple trace*) of the foraging model are shown, along with the objective reward probabilities (*pale traces*). The horizontal green line is the threshold. **C)** Distribution of the probability of choosing the objectively best option for both foraging (*green*) and RL (gold) agents. **D)** Difference between the distributions seen in *C*. **E)** Distribution of the normalized average reward received by both foraging (*green*) and RL (*gold*) agents. **F)** Difference between the distributions seen in *E* (caret = mean). **G)** the negative log-likelihood of the RL and foraging models fit to all data from Experiment 1. **H)** The Akaike information criteria (AIC) of the RL and foraging model fits. **I)** Average choice likelihood for each participant under the foraging model (x-axis) and the RL model (y-axis). *Inset* : Distribution of differences in choice likelihood across participants of Experiment 1. **J)** The negative log-likelihood of various extensions of the RL (*gold*) and foraging models (*green*). **K)** The Akaike information criteria of the extensions of the RL (*gold*) and foraging models (*green*).

Simulating both agents in the same reward schedules experienced by our participants revealed that the environment did not advantage foraging over RL; in fact, RL agents outperformed foraging agents, regardless of whether performance was defined as the probability of choosing the objectively best option (Wilcoxon signed rank test : *p <* 0.01; Figure 3C/D) or as the probability of reward (Wilcoxon signed rank test : *p <* 0.01; Figure 3 E/F). In short, the task did not encourage compare-to-threshold decision-making and, in fact, the participants would have had to sacrifice reward to adopt this strategy over a compare-alternatives strategy.

### Foraging models better predict participant decisions

To determine how the participants solved the task, we fit computational models (see Methods). We first compared a simple 2-parameter Reinforcement learning model (“standard RL”; Rescorla-Wagner; [1, 6]) with a simple 3-parameter formulation of the foraging model (“standard F”). We found that the standard foraging model was a better fit to participant behavior than the standard RL model (Figure 3G-H; standard F: log-likelihood = -23,206, AIC = 47,936, 3 parameters by 254 participants = 762 parameters; standard RL: log-likelihood = -24,878, AIC = 50,771, relative AIC weight *<* 1.8*10^*−*32^, 2 × 254 participants = 508 parameters; 76,200 total trials). The foraging model also outperformed the RL model on an individual basis: it was a better fit in 64.2% of individual participants (163/254; Figure 3I; standard F: median individual log-likelihood = 82.36, 95%CI = 9.40 to 201.06, standard RL: median = 86.72, 95% CI = 9.56 to 199.66; significant paired t-test: *p <* 0.0001, t(253) = 6.79, mean difference = 5.27, 95% CI = 3.74 to 6.79). This suggests that a foraging-like compare-to-threshold mechanism may better describe the participants’ choices in this task.

There are a variety of extensions to the RL model that outperform the standard 2-parameter formulation and it remained unclear whether any of these would fit behavior better than foraging. Therefore, we next compared the foraging model with 4 common extensions of the RL model (2 forms of choice history dependence, asymmetrical learning, and value decay, see Methods). We found that every extended RL model performed substantially better than the standard RL model in model comparison (Figure 3J-K). However, the foraging model continued to outperform: it was a better fit than RL models that incorporated asymmetrical learning from wins and losses (“asymmetrical RL” [24, 29, 30]; log-likelihood = -23,963, AIC = 49,450, AIC weight relative to foraging model *<* 10^*−*32^, 762 parameters), decay in the value of unchosen options (“decaying RL”; [15, 31]; log-likelihood = -23,181, AIC = 47,887, relative AIC weight *<* 10^*−*32^, 762 parameters), and a simple form of choice-history dependence with 1 added parameter (“history-kernel 1 RL”; [4]; log-likelihood = -23,252, AIC = 48,028, AIC weight relative to foraging model *<* 10^*−*32^, 762 parameters). The only extended RL model that outperformed the simple foraging model was a 2-parameter choice-history dependent model (“history-kernel 2 RL”; [4]; log-likelihood = -22,660, AIC = 47,351, relative weight of the foraging model *<* 10^*−*32^). Thus, the simplest foraging model was a better explanation for behavior than all but the most complex of RL models.

The various extensions of the RL models were developed over a period of decades to improve model fit and it remained ambiguous whether the 1 RL (*i*.*e*., 2-parameter choice-history dependent RL) model that did better than the foraging model did so because it was a compare-alternatives model or because it incorporated history dependence. Therefore, we next developed an equivalent version of the foraging model for each extensions of the RL model and compared the two model classes (see Methods; Figure 3J-K). Each extended foraging model outperformed the equivalent RL model. This was true for a foraging model that permitted asymmetrical learning between wins and losses (“asymmetrical foraging”: log-likelihood = -22,777, AIC = 47,585, AIC weight relative to asymmetrical RL *<* 10^*−*32^, 4 × 254 = 1016 parameters), for a foraging model where the threshold decayed with repeated choices (“decaying foraging”: log-likelihood = -22,632, AIC = 47,295, AIC weight relative to decaying RL *<* 10^*−*32^, 1016 parameters), and for the 1-parameter history-dependent model (“history-kernel 1 foraging”: log-likelihood = -22,558, AIC = 47, 148, AIC weight relative to choice-kernel RL 1 *<* 10^*−*32^, 1016 parameters). Ultimately, the best model overall was the history-dependent foraging model, which outperformed the best RL model (“history-kernel 2 foraging”: log-likelihood = -22,292, AIC = 47,125, AIC weight relative to choice-kernel RL 2 *<* 10^*−*32^, 1270 parameters; relative AIC weights of all competing models *<* 10^*−*32^). This foraging model also outperformed the best RL model on an individual basis: it did at least as well or better in nearly every individual participant than the best RL model (foraging model: median individual log-likelihood = 79.32, 95% CI = 8.92 to 197.18, RL model: median = 80.43, 95% CI = 8.72 to 195.31; significant paired t-test: *p <* 0.05, t(253) = 1.98, mean difference = 1.45, 95% CI = 0.01 to 2.89). In short, as a class, the foraging models consistently provided a better fit to the participants’ decisions.

It remained possible that the foraging model outperformed the RL model solely because it was a better fit for participants who did not fully understand or engaged with the task. However, there was no systematic relationship between how foraging-like an individual participant was and how well they did in the task. Individual differences in the “foraging index” (see Methods) were not correlated with the probability that the participant would choose the best option (*R* = −0.01, *p* = 0.84) nor were they correlated with above-chance reward probability (*R* = 0.03, *p* = 0.64). In sum, the participants who were most likely to be using foraging computations did not have either an advantage or a disadvantage in this task. Considering that foraging agents significantly underperformed RL agents (Figure 3C-F), this null result suggests that our most foraging-like participants were just as engaged in the task, if not moreso, than other participants.

### Foraging better explained the participants’ tendency to repeat

We next asked why the foraging model was a better fit to behavior: what aspects of behavior did it describe that the RL model was unable to account for? We previously found that choices are considerably more repetitive than what can be explained with standard RL models, at least in humans and other primates [15, 32]. This may explain why most common extensions of the RL model tend to make the model more repetitive. This is most obvious in the case of the choice history kernel models: the added parameters increase the probability of repeating the previous choice. In the decay model, similarly, repetition increases as the value of the alternative decays towards zero. Even in the asymmetrical learning models, learning more from wins than losses tends to stabilize choices and increase repetition [33, 34]. Therefore, we next asked if the foraging model was a better fit to behavior because it also better captured the repetitiveness of the participants’ choices.

We simulated data from RL and foraging agents that were matched to the participants’ environment and parameters (see Methods). We then compared the level of repetitiveness in these simulated datasets with the participants’. In the participants, the average run length (length of same-choice sequences: 4.31 trials) was close to our expectation under foraging (4.33 trials, 95% CI = [4.24, 4.43]) but greater than any individual run length from the RL simulations (4.18 trials, 95% CI = [4.09, 4.24]; *p <* 0.01; Figure 4A). This was related to the fact that the foraging model was more accurate in predicting how often the participants would switch (i.e., the inverse of the choice run lengths; Figure 4B-D; foraging, mean sum of squared errors [MSSE] = 0.72, 95% CI across simulations = [0.53, 0.94], RL, MSSE = 2.78 (95% CI = [2.54,3.01]; significantly greater in RL; 2-sided paired t-test: *p <* 0.0001, t(1,253) = 6.46). The average probability of switching under the RL model was 22.7% (95% CI = [22.3%, 23.1%]) and under the foraging model it was 20.3% (95% CI = [20.0%, 20.7%]). The participants switched 20.0% of the time, less frequently than any of the samples from the RL model, but not the foraging model. In fact, the RL model systematically predicted that individual participants should switch more than they did (2.8% more, 95% CI across individuals = [1.7%, 4.0%], *p <* 0.0001, t(1,253) = 4.72). The foraging model did not have a significant bias towards over- or under-estimating individual participants (predicted 0.4% more on average, 95% CI = [-0.08% to 0.1%], p = 0.1, t(253) = 1.66). In short, in comparison to the RL model, the foraging model better predicted the participants’ tendency to switch on both the individual and group levels: it was both more precise and less biased.

**Fig. 4:**
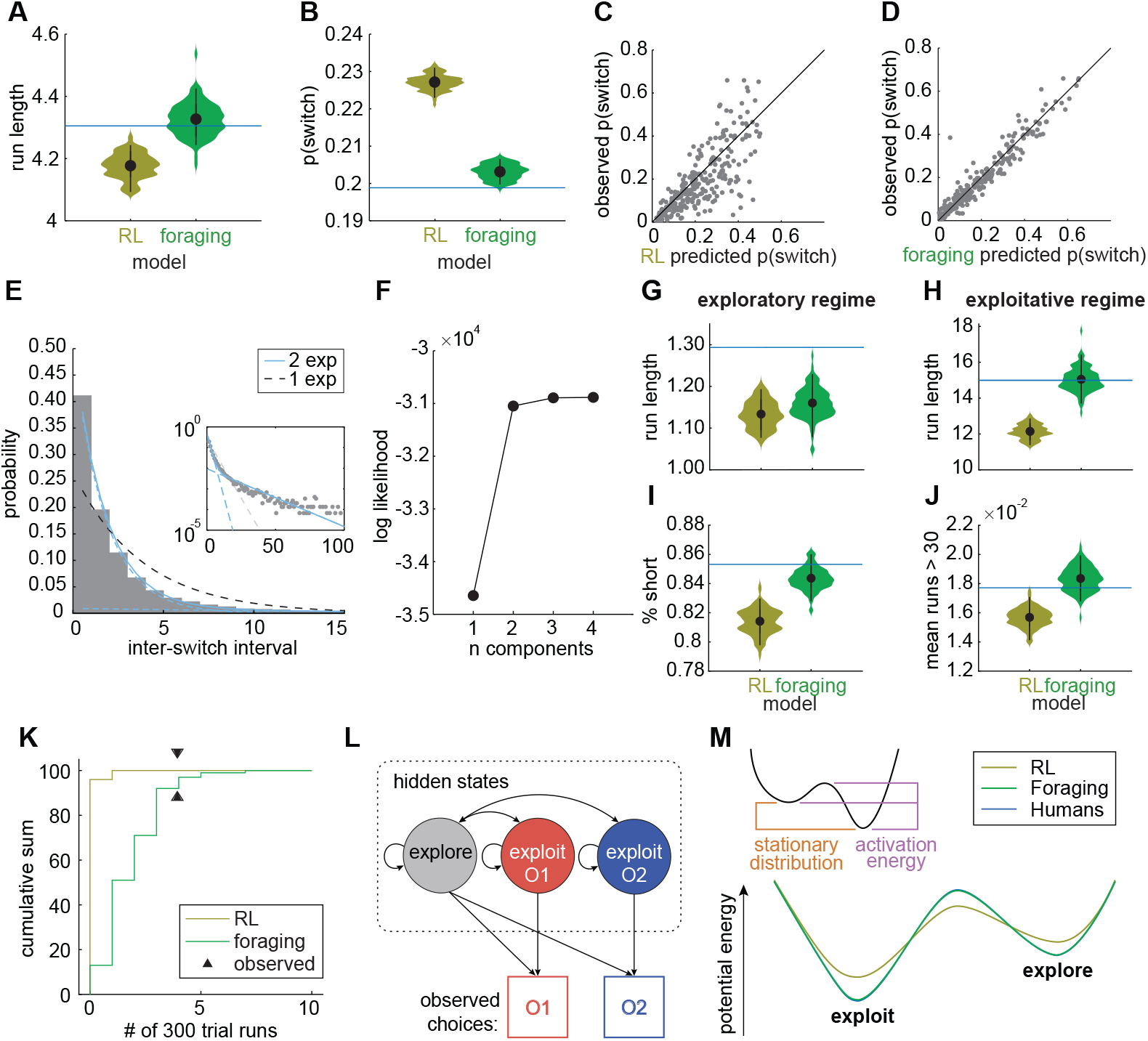
Foraging and RL predict different choice dynamics. **A)** Average number of trials between switch decisions (*i*.*e*., run length) of RL (*gold*) and foraging (*green*) agents in simulated Experiment 1 (blue line = average for Experiment 1 participants). **B)** Probability of switch. Same convention as in *A*. **C)** Human observed probability of switch as a function of RL predicted probability of switch. **D)** Human observed probability of switch as a function of foraging predicted probability of switch. **E)** Distribution of choice run lengths of humans in Experiment 1. If the probability of switching was fixed, run lengths would be exponentially distributed (*black dotted line*). A mixture of two exponential distributions (blue line) suggests 2 distinct probabilities of switching. Dotted blue lines show each mixing distribution, one slow-switching (*i*.*e*., exploitative regime) and another fast-switching (*i*.*e*., exploratory regime). *Insert*: Same data presented on a logarithmic scale. **F)** Log-likelihoods for different mixture models containing a range of 1 to 4 exponential distributions. **G)** Average run length in the exploratory regime. **H)** Average run length in the exploitative regime. **I)** Proportion of exploratory regimes. **J)** Proportion of very long exploitative regimes exceeding 30 consecutive trials. **K)** Four participants were held out of all analyses because they chose the same option 300 trials in a row. To determine the expected number of these no-switch participants under each model, 100 datasets were generated from the distribution of fitted parameters (each with 254 simulated participants). Here, we plot the cumulative percent of simulations (y-axis) as a function of the numbers of no-switch participants (x-axis). Black caret = proportion of no-switch participants in Experiment 1. **L)** Hidden Markov model (HMM) was used to infer the underlying internal state of the participants on each trial from their sequence of choices. The model included 3 hidden states, 2 exploitative states corresponding to each option and an exploratory state which participants could choose any of the options. **M)** Overlaid state dynamic landscapes for both foraging (*green*), RL (*gold*) agents and Experiment 1 participants (*blue*).

### Foraging better explained persistent choice runs

In mice, monkeys, humans, and optimized RL models, the distribution of choice run lengths in this task is composed of two switching regimes: one regime with rapid switching that is likely due to exploratory trial-and-error sampling and one regime with slow switching that is likely due to exploitative, persistent choices to the same target [15, 24, 32]. A specific increase in the slow switching regime is what sets humans and other primates apart from rodents in this task [32] and it is also one major feature of human decision-making that RL models do not naturally capture [15]. Therefore, we next asked whether the foraging model might better account for the slow switching regime.

Again in this dataset, the behavior of the participants and both models were well described as a mixture of 2 regimes (Figure 4E-F; see Exponential Mixture Models in Methods). In the participants, the 2 regime model (log-likelihood = -31,048, 3 parameters, n = 14,847) provided a significantly better fit than 1 (log-likelihood = -34,645, 1 parameter, likelihood ratio test: ratio = 7193.0, p *<* 10^*−*32^) and, while adding additional switching regimes continued to improve model fit (3 regimes: log-likehood = -30,893, 5 parameters; 4 regimes: log-likelihood = -30,884), improvement was already saturated at 2 (Figure 4F). Two regimes were also apparent across all the simulated data from the RL and foraging models (RL: n = 1,699,789, 1 regime log-likelihood = -3,907,700, 2 regimes log likelihood = -3,527,300, 3 regimes log-likelihood = -3,502,000, 4 regimes log-likelihood = -3,500,600, significant improvement from 1 to 2: ratio = 760,950, p *<* 10^*−*32^, elbow at 2; foraging: n = 1,517,656, 1 regime log-likelihood = -3,550,000, 2 regimes log likelihood = -3,128,100, 3 regimes log-likelihood = -3,105,700, 4 regimes log-likelihood = -3,103,900, significant improvement from 1 to 2: ratio = 843,650, p *<* 10^*−*32^, elbow at 2). In sum, both models and the participants had two distinct switching regimes in this task.

The switching regimes in the foraging model better matched the participants than those in the RL model. Both the foraging and the RL model tended to switch more frequently than the humans did during their own fast-switching, exploratory regimes (Figure 4G; RL = 1.13 trials, 95% CI = [1.08, 1.19]; foraging = 1.16, 95% CI = [1.08,1.23]; participants = 1.29, significantly longer than both models, *p <* 0.01). This may be due to the fact that neither model accounted for the complexity of exploratory decision-making, which can be information maximizing rather than random in some circumstances [35–37]. Nonetheless, only the foraging model was able to replicate the average long run length of the participants’ slow-switching, exploitative regimes (Figure 4H; RL = 12.15 trials, 95% CI = [11.38, 12.86]; foraging = 15.05, 95% CI = [13.70,16.42]; participants = 14.99, only significantly different than RL [*p <* 0.01]) and their relative frequency (Figure 4J). This might explain why participants had more very long choice runs than we would expect under RL (1.8% of their choice runs were longer than 10% of the total number of trials [30 trials], significantly more than the 1.6% predicted by RL, 95% CI = [1.4%, 1.7%], *p <* 0.01) (Figure 4J). The participants’ long-run lengths frequency was well within the distribution under the foraging model, however (1.8%, 95% = [1.7%, 2.0%]). In sum, the foraging model better accounted for the participants tendency to repeatedly persist in choosing certain options for long periods of time.

Recall that we initially excluded 4 of our 258 participants (1.6%) because they chose same option for the entire 300-trial duration of the session, despite passing our initial screening criteria. Because model parameters were not identifiable in these participants, they represented held-out data that did not influence the simulations in any way. Therefore, we next asked how likely these participants were, given the two models. In the simulated data, we found that runs of 300 identical choices were very rare in RL simulations (4 of 25,400 or 0.016% of simulated sessions), in contrast to foraging (178 of 25,400 or 0.7% of simulated sessions). This meant that 4 or more participants who chose the same option 300 trials in a row would be expected in 10.9% of experiments under foraging (Figure 4K). By contrast, under RL, we would have less than a 1 in 1 million chance of observing these 4 participants. In sum, the foraging model better captured the stability or repetitiveness of participants choice runs, in both those that were included in the model fits and in those that were excluded.

### Foraging better explains the dynamics of switching and exploitation

Foraging was better able to explain the participants’ tendency to generate long choice runs. This could suggest that the foraging model was simply more stable than the traditional RL model. However, the foraging model also predicted that participants should have more fast-switching (*i*.*e*., exploratory regime) choices than the RL model did (Figure 4B/I; relative frequency of short choice runs; foraging = 84.4%, 95% CI = [82.8%, 86.0%]; RL = 81.4%, 95% CI = [79.8%, 83.0%]). Again the participants matched the predictions of the foraging model, but not the RL model: they had many more short choice runs than the RL model had predicted (relative frequency of exploitative runs = 85.3%, only significantly different than RL [*p <* 0.01]). In short, the participants both switched more and switched less than the RL model suggested they should, but foraging was able to capture the complexity of the switching dynamics.

To understand why the foraging model was better at reproducing the participants’ switching dynamics, we used latent state models to interrogate the dynamics of exploration and exploitation in the participants and in both models (see Methods; [15, 24, 36, 37]. Briefly, this approach models exploration and exploitation as the latent states underlying sequences of decisions through training hidden Markov models (HMMs) on observed choice sequences (Figure 4L). Where the mixture models helped characterize the switching dynamics from runs of sequential choices, the HMMs were used to infer the most likely generative state underlying each individual choice [15, 24, 36] and to make statistical inferences about the governing equations of exploration and exploitation [24, 32, 36]. For example, the parameters of the transition matrix of an HMM are estimates of the probability of transitioning between specific states: that a decision-maker will continue to exploit once they have begun to exploit, for example, or that they will stop exploiting one option in order to explore the alternatives.

The foraging model better matched the participants’ explore/exploit state dynamics than did the RL model. Every measurement made with the HMMs was out of sample for the RL model simulations, but well within sample of the foraging model. For example, the participants repeated exploration 90.4% of the time, which was significantly higher than the RL model simulations (87.3%, 95% CI = [86.3%, 88.4%], *p <* 0.01) but close to the mean of the foraging simulations (90.3%, 95% CI = [89.8%, 90.9%]). Similarly, the participants repeated exploitation 93.9% of the time, which was significantly higher than RL (90.9%, 95% CI = [90.5%, 91.3%], *p <* 0.01) but close to the mean foraging model (93.7%, 95% CI = [93.4%, 94.1%]). The fact that the transition matrix of an HMMs is a Markov chain makes it particularly analytically tractable. We can often solve for the stationary distributions of these equations: the long-term probability that the participants would exploit (vs. explore; see Methods). Here again, we found that the participants stationary frequency of exploration and exploitation (39.2% explore, 60.8% exploit) was out of sample for the RL model (41.6% explore, 95% CI = [40.4%, 42.9%]; 58.4% exploit, 95% CI = [57.1%, 59.6%], *p <* 0.01 in both cases), but well within sample for the foraging model (39.3% explore, 95% CI = [38.4%, 40.2%]; 60.7% exploit, 95% CI = [59.9%, 61.6%]). Together, these results illustrate that the foraging model well described the dynamics of exploration and exploitation in the participants, while the RL model was consistently off target.

Plotting the energetic landscape of these dynamics (see Methods; [24, 32, 36]) revealed the intuition behind all of these individual results: the energetic landscape of exploration and exploitation was flatter in the RL model than it was in either the participants or the foraging model (Figure 4M). In both the participants and the foraging model, exploitation was a deeper, more stable state than exploration, but a substantive energetic barrier between the two states meant that exploration and exploitation were actually fairly stable states, with infrequent transitions between them. Conversely, in the RL model, not only was there less of a difference in the depth of exploration and exploitation, but there was less of an energy barrier between. Together, these results suggest that the foraging model was a better fit for the participants because it better captured the dynamics of exploration and exploitation: their tendency to alternate between temporally-extended periods of exploiting good options and exploring alternatives.

### Foraging better predicts behavior under varied uncertainty

It is possible that the participants did not bother comparing alternative values because the value of unchosen options was difficult to estimate in a restless bandit environment. Therefore, in Experiment 2 and 3, we bidirectionally manipulated the uncertainty of the unchosen option to to try to make the participants more or less likely to prefer the RL-like, compare-alternative computations over the foraging-like compare-to-threshold.

In Experiment 2, participants (95 people, 41 females, 1 other or non reporting) performed an more predictable version of the 2-armed bandit task where information about the unchosen option was increased. In this variation, the environment was structured so that the sum of reward probabilities for any given trial always totaled one (‘Example walk’, Figure 5A). This reduces the uncertainty of the unchosen option because its value could be inferred from the chosen option (it was always 1 minus the value of the chosen option). In this new task, we found that foraging was also a better fit to human behavior than RL (Figure 5B-C, FOR: log-likelihood = -6819, RL: loglikelihood = -7241 and relative AIC weight *<* 5.1 *10^*−*32^). (Note that we were unable to fit the model in 1/95 participants because that participant repeatedly selected the same option 300 times ([1.1%]; a similar proportion to the number that did so in Experiment 1: 4/258 [1.5%]).) To understand why foraging was a better fit, we generated an RL and foraging agent from each participant’s parameters (see Methods) and then compared the simulated agents’ behaviors with the humans’ choice patterns. In line with Experiment 1, foraging better accounted for the participants’ tendency to switch between options (Figure 5D-E, RL: MSSE = 0.38, foraging: MSSE = 0.06, Wilcoxon signed rank test *p <* 1.5 * 10^*−*7^). Together, these results suggest that foraging continues to be a better fit even when it is easier to infer the value of the unchosen option.

**Fig. 5:**
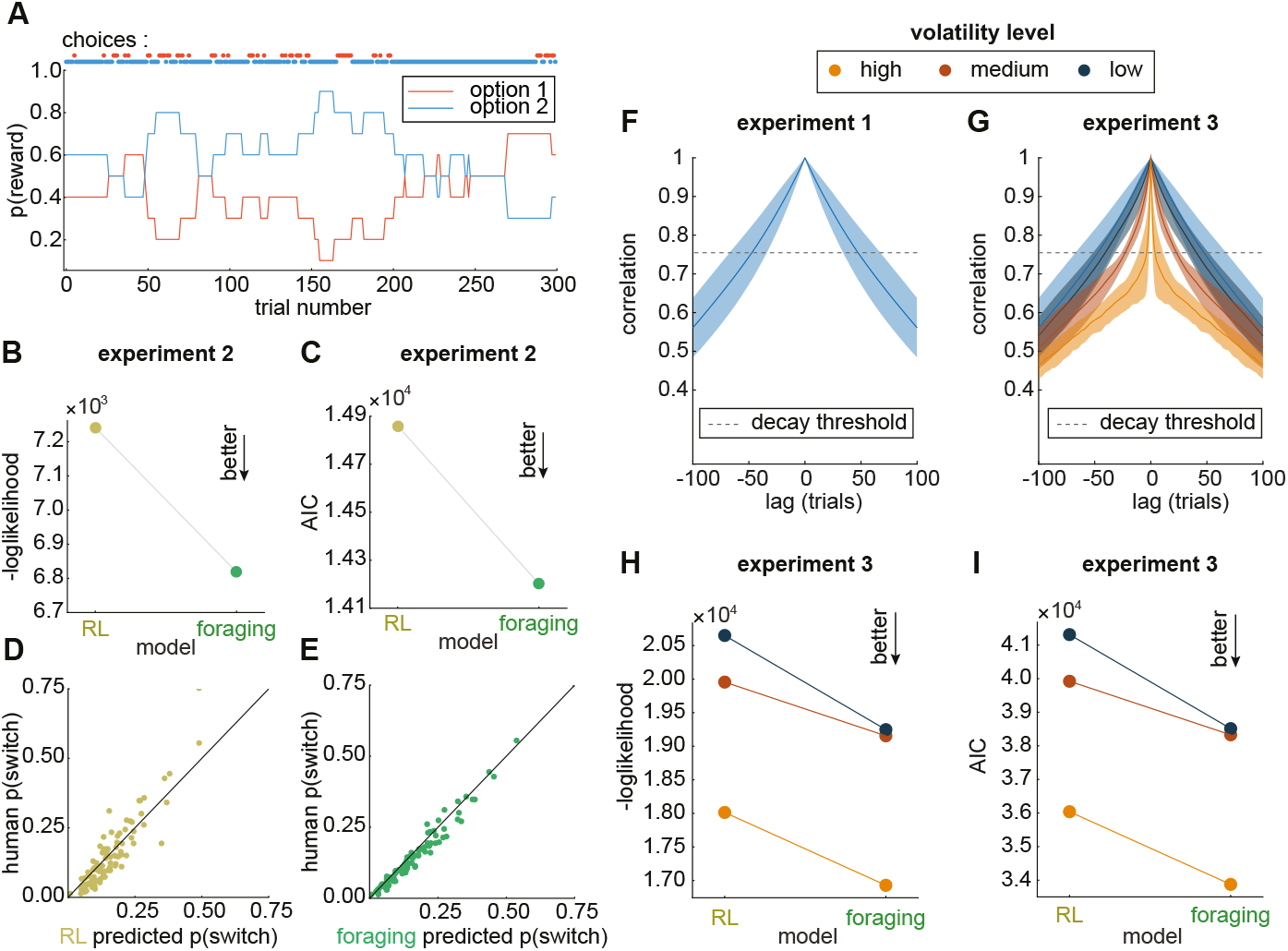
Foraging and RL have different quality fit to the data across different environments. **A)** An example reward schedule from the predictable two-armed bandit task (Experiment 2), illustrating the reward probability of each option (red trace = option 1; blue trace = option 2) and the choices made by the participant who had this reward schedule (*red dots* = option 1;*blue dots* = option 2). **B)** The negative log-likelihood of the RL and foraging models fit to all data of Experiment 2. **C)** Akaike information criterion. Same convention as in *B*. **D)** Human observed probability of switch in Experiment 2 as a function of RL predicted probability of switch. **E)** Foraging predicted probability of switch. Same convention as in *D*. **F)** Reward schedules auto-correlations averaged for all participants of Experiment 1. Full line indicates the mean, shading indicates STD and dotted line indicate the decay threshold. **G)** Same convention as in *F* for 3 levels of volatilities (*red line*) participants of experiment 3 **H)** The negative log-likelihood of the RL and foraging models fit to all data of Experiment 3. **I)** AIC of the RL and foraging models across different volatility levels in Experiment 3. Same convention as in *G*.

In Experiment 3, participants (270 people, 112 females, 2 other or non reporting) performed a 3-armed version of the bandit task in which information about the unchosen option was reduced by (1) manipulating volatility and (2) increasing the number of options (see Methods). Increasing the volatility of the environment increases uncertainty about unchosen options because it increases the speed at which prior reward information becomes uninformative about the current value. Moreover, prior research suggest that increasing volatility tends to increase switching behaviors in humans [38, 39], which could offer an advantage to RL, as a less stable and persistent algorithm. Given that both Experiment 1 and 2 had similar low volatility (Experiment 1 is illustrated in Figure 5F), we increased the volatility of the environment by low, medium, or high levels (Figure 5G). Regardless of the volatility of the environment, foraging was a better fit (Figure 5H-I, “high volatility” FOR: log-likelihood = -16,928, RL:log-likelihood=-18,013 and relative AIC weight *<* 1.2 * 10^*−*32^; “medium volatility” FOR: log-likelihood = -19,155, RL:log-likelihood=-19,955 and relative AIC weight *<* 5.5 *10^*−*32^; “low volatility” FOR: log-likelihood = -19,247, RL:log-likelihood=-20,649 and relative AIC weight *<* 1.8 * 10^*−*32^). Together, these results suggest that foraging better explained human behavior even in high volatility environments that should have been best suited to the kind of frequent switching seen in RL models.

## Discussion

Although human and animal behavior in k-armed bandits are conventionally modeled as a compare-alternatives process [13–15, 24, 28], our results suggest that humans compare the value of staying with the currently exploited option against a fixed threshold rather than comparing the values of choosing alternative options directly. We leveraged this finding to build a novel compare-to-threshold RL model (“foraging”) and compare it to a traditional compare-alternatives RL models [1, 4, 6]. The foraging model and its various extensions were a better fit to participants’ behavior compared to the compare-alternatives RL models [1, 4, 6] even when comparing alternatives would have been the more advantageous strategy. The foraging model fits were consistently better than traditional RL models across 2 variations of the task that manipulated the uncertainty of the unchosen option. Across environments, the foraging models consistently better matched participants’ tendency to switch options, both on average and on an individual basis. The foraging models were even able to predict the existence of multiple participants who never switched at all: participants that essentially should not have existed under compare-alternatives RL.

The difference between the foraging and traditional RL models is not solely that the latter uses compare-alternatives computations and the format uses compare-to-threshold. Another critical difference is in the models’ action-value space. In the traditional RL model, values represent the subjective utility of each primitive action (choosing option 1 or 2). By contrast, the foraging model does not represent the value of primitive actions, but instead of temporally extended actions (*i*.*e*., of exploiting the left option, rather than choosing the left option). In this way, the foraging model resonates with recent work on hierarchical reinforcement learning that suggests that people do make choices at the level of policies rather than primitive actions [40–46]. Higher level policies, macro actions and options [40] have been studied for over two decades in the RL literature and are a candidate strategy for achieving better generalization and transfer of knowledge in artificial agents [40, 43]. This distinction between models could make the foraging model better able to scale outside of laboratory tasks, where the number of options available is generally much larger than the small finite sets we present in the lab. Where the computations and representations in traditional RL models become more costly in high-dimensional or continuous decision spaces, the computations and representations in the foraging model can remain comparatively efficient. Compare-to-threshold computations may thus be favored by humans because they better align with the brain’s biological constraints and because they are better equipped to scale to natural environments. Together, these considerations suggest that foraging is a better explanatory model both because decisions are more like compare-to-threshold than compare-alternatives, but also because decisions are made at the level of temporally extended policies, rather than primitive actions.

The foraging model we introduce here draws both its inspiration and its limitations from the foraging literature and could be expanded on in the future in ways that could have implications across fields. The model assumes the existence of only two macro actions—explore and exploit—and defines a specific low-level policy and termination for each of them. Based on the foraging literature, we assumed that switching from exploit to explore causes instant forgetting of exploited values (i.e. the patch is left behind), that explore actions select targets randomly (i.e. that we travel until we encounter some new opportunity at random), and that exploit action-values are compared to a fixed threshold (i.e. one that is learned very slowly across long-term experience in an environment). On one hand, drawing these assumptions from the ethological literature is a powerful way to create new computational models and constrain the space of useful macro polices. On the other hand, each of these assumptions is also an opportunity for future research that could improve on the simple foraging algorithm presented here. For example, the foraging model failed to account for the rapid speed of switching during the exploratory regime(Figure 4G). This might suggest that refining the exploratory policy in this model could substantially improve both model fit and agent performance. One promising approach would be to incorporate directed forms of exploration (like self-avoiding search [37]) or intrinsic motivations (like entropy maximizing policies [47]). Developing more sophisticated exploratory strategies is likely necessary for translating the simple foraging algorithm we present here into something that could be useful in AI. In that sense, our work could represent a bridge between observations in animal behavior and major challenges in AI, including the problem of identifying useful macro action spaces. In short, developing new, biologically-inspired models of human decision-making could have broader impacts: not only for understanding the human mind, but also for generating powerful new tools in AI.

The observation that foraging models are a better fit to human behavior may superficially appear to contradict the large body of evidence for RL-like computations in the neuroscience literature. However, the neural evidence for reinforcement learning computations in the brain is strongest when we consider signals related to *reward prediction errors* and *value functions*— both of which also exist in our novel foraging model. Both models calculate reward prediction errors: this is how they learn their respective value functions from their environments. Further, although neural activity often correlates with action value (a signal thought to play a role in specifying primitive actions; [48]), RL primitive action-values are likely to be highly collinear with the macro action-values calculated by the foraging model. Because no direct comparison between these two value signals has been performed, it is thus not entirely clear that evidence for the former would be evidence against the latter. We would note that it is always important to be cautious in interpreting correlations between value and neural activity because (1) time series data are prone to spurious correlations, (2) putative neural correlates of value could be caused by other mechanisms, like mnemnonic errors [49], and (3) neural activity have a complex, nonlinear relationship with value even when value is explicitly cued [50], which calls into question the interpretability of linear correlations here. Further, there exist some neural observations—such as nonlinearities in neural activity around shifts between exploration and exploitation [15, 25, 51–53]—that are difficult to reconcile with traditional RL models. In the foraging model, by contrast, instant forgetting at the onset of exploration causes an obvious nonlinearity. It will be important, in the future, to determine whether these assumptions of the forging model are either validated or refuted by neural observations [15, 25]. Ultimately, neuroscience research will hold the key to determining whether the computations humans perform are truly more consistent with compare-alternative or compare-to-threshold accounts.

Ultimately, this paper reinforces and extends a body of recent observations that humans, like many other animals, can use foraging computations to make sequential decisions [19–21]. However, where previous studies of foraging computations in psychology and neuroscience used foraging-specific tasks—tasks that are explicitly designed to encourage foraging strategies or computations [18–23, 54, 55]—the present study did the opposite. Through developing a new foraging model that is capable of navigating dynamic, unpredictable environments, we determined that humans also use foraging computations even in tasks that were originally designed as testbeds for compare-alternatives RL algorithms.

## Methods

### Data Collection

#### Ethics Approval

All experiments procedures were in line with the standards set by the Declaration of Helsinki and were approved by the relevant ethical review boards (Experiment 1 and 2: the guidelines of the Comité d’Éthique de la Recherche en Sciences et Santé (CERSES) of the University of Montreal [Project #2021-1090], Experiment 3: Institutional Review Board of the University of Minnesota [STUDY00008486]).

#### Participants

Participants were recruited via the online platform, Amazon Mechanical (mTurk). To avoid bots and facilitate better data quality, participants were only accepted when they had a minimum of 5000 approved human intelligence tasks (HIT) and a minimal percentage of 98% in proportions of completed tasks that are approved by requesters. Geographical restrictions were set for US participants only. To help ensure that participants understood the task, there was an initial 25 trial block with fixed but randomly assigned reward probabilities (20%, 30% and 70%). To continue to the main task, participants were required to (1) switch between options at least twice, and (2) get reward more than 42% of the time during the practice block. The criteria and number of trials was chosen to ensure that people choosing at random would rarely pass through to to the main task. Participants were not allowed to repeat the experiment. All participants who successfully submitted the HIT, either the practice block or the full experiment, were paid a base rate of $0.50, plus a bonus of $3.61 mean ± $0.97 SD based on their performance (for each trial that ended with a reward, participants were given a $0.02 compensation). Participants provided written informed consent after the experimental procedure had been fully explained and were reminded of their right to withdraw at any time during the study.

For Experiment 1 (two-armed bandit task), a total of 258 participants (120 females, 137 males, 1 preferred not to say) completed the task. This dataset has been analyzed previously in a comparison of switching behavior between mice, monkeys, and humans [32], but all analyses presented here are new. For experiment 2 (predictable two-armed task), a total of 95 participants (41 females, 51 males) completed the task. For experiment 3 (three-armed bandit task), a total of 270 participants (112 females, 156 males, 2 prefer not say) completed the task.

#### Experiment 1: Two-armed restless bandit task

To investigated how humans solved a restless bandit task, participants were required to repeatedly choose one of two options. Each option was associated with a probability of reward, which changed across trials and independently across options. The experiment consisted of 25 practice trials followed by 300 trials. To avoid environmental biases, each participant’s reward schedule was randomly initialized. At any given step, the reward probability of each option could subsequently vary by a 0.1 increment with a hazard rate of 0.1, following a Bernoulli distribution.

#### Experiment 2: Predictable 2-armed bandit task

To understand how decreased uncertainty in unchosen option influences behavior, participants performed a modified version of the two-armed restless bandit task. Although both tasks are similar, they differ in how rewards are allocated to each of the two options. Specifically, in this task, the reward probabilities associated with each option varied inversely with respect to one another and across time.

#### Experiment 3: Volatile 3-armed bandit task

To explore how increased uncertainty in unchosen options influences behavior, participants performed a three-armed restless bandit task under various manipulations of volatility of the environment. This task differed from the two-armed restless bandit in two keys aspects. First, an additional option was introduced. This is common practice in the field, and previous studies show that increasing the number of options (up to 4) has no significant effect on the probabilities of switching and exploring [32]. Second, the change in reward probabilities was variable (probability of step = 1 or 0.1), with a different increment size for each participant (ranging from 0.05 to 0.667).

Since there were 2 manipulations in this task (i.e., probability of step and step size), volatility was assessed by looking at the auto-correlation of the reward schedule experienced by each participant. We estimated the rate of decay of the auto-correlation by measuring when it decayed to 0.75 of its initial value. We used this measurement as an arbitrary volatility index for each participant. We then divided the volatilities into 3 levels (high, medium, low).

## Data Analysis

### General Analysis Methods

Data was analyzed with custom software written in MATLAB and Python. Statistical tests were two-sided unless otherwise specified. Significance was compared against the standard alpha = 0.05 threshold throughout. Correlations are Pearson correlation unless otherwise indicated. A small number of participants (Experiment 1: 4/258 participants, 1.6%; Experiment 2: 1/95, 0.95%) were excluded from some model-based analyses because they chose only one option in the main task, which made the model parameters unidentifiable. However, these participants were included in other analyses wherever it was possible to do so.

### Strategy Fingerprints

Compare-alternatives and compare-to-threshold decision-making depend on different aspects of the environment. Here, we characterized the fingerprints of each strategy as a function of two features of the environment: richness (the sum of the values) and discriminability (the range of the values, normalized to the sum). We assumed that the compare-alternatives strategy would switch whenever discriminability was below some Δ reward and that the compare-to-threshold strategy would switch whenever richness was below some threshold. Example simulations for specific parameter values of Δ reward and threshold are illustrated in 2B-C. To quantify the fingerprints, we fit the simulated switch probabilities at each parameter value for each strategy with a linear (2 parameter) generalized linear model (GLM) and a quadratic (3 parameter) GLM, first as function of richness and then as a function of discriminability. (Identical results were found with the identity and binomial link functions, though only the former was used for illustration.) We then extracted 3 features from the fitted models: (1) the relative deviance of the linear and quadratic models (a measure of the extent of curvature as a function of either richness or discriminability), (2) the quadratic term of quadratic model (a measure of the sign and magnitude of curvature in the relationship), and (3) the linear term of the linear model (a measure of the overall slope of the relationship). Distributions of these features are illustrated in 2E-G. To provide a more holistic and multivariate view of these fingerprints, we took the 6 feature dimensions described above and added the overall probability of switching as a 7th feature. We then performed PCA for illustration (2H) and calculated the distance from the 7-dimensional fingerprint of the human data to the fingerprint of every possible value for each of the two strategies 2I.

### Foundations of Decision-Making Models

To determine whether participants were using foraging-like or RL-like computations, we fit a series of 10 cognitive models (5 RL models and 5 foraging models). All models were initialized with 50 random seeds and fit via maximum likelihood (minimizing the negative of the log likelihood; fminsearch, MATLAB, scipy.optimize.minimize, Python). Here, we will describe the standard formulations of both the RL and foraging models. The following section will explain the various extensions of each model.

All of the models, RL and foraging alike, had a classic delta-rule value updating process at their core,

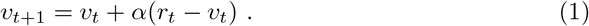

That is, values (*v*_*t*_ +1) were updated at each time step *t* according to the difference between the observed reward (*r*_*t*_) and the previous value (*v*_*t*_; *v*_0_ = 0 for all values and in all models). The magnitude of the update is scaled by a learning rate (*α*; constrained between 0 and 1 in all models). The delta-rule update dynamically calculates value as an exponentially-decaying recency-weighted average of reward. This is a canonical computation, with widespread neural and behavioral evidence supporting its existence in humans and other animals [1, 6, 14, 56–60]. Though both models have the delta-rule computation at their core, they differ in what the value represents.

#### RL model

Here, there are multiple values because each option *i* has its own value and each is updated (or not) at each time step. One common approach, which we used in the standard RL model here, is to update only the value of chosen options,

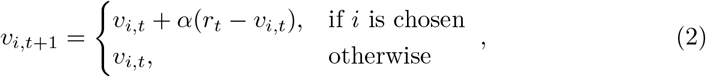

A softmax function transformed values into choice probabilities with the usual Boltzmann exploration strategy.In RL models, the probability of choosing option *i* at time *t* was

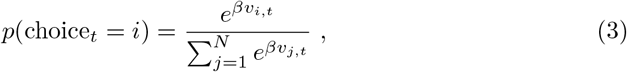

where *β* is an inverse temperature parameter that controls the noise in the value comparison (constrained between 0 [high noise] and 100 [low noise] in all models). Because this task only had *N* = 2 options, we can expand the sum and rearrange this equation,

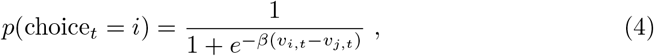

which may clarify the point that this function compares the value of the two options in order to decide which to choose.

#### Foraging model

Here, the value is the value of a single focal option: it is the value of staying with or “exploiting” that option, so the update equation is precisely Eq. 1. One useful way to characterize the foraging model is to think of it as a hierarchical action algorithm: there are higher-level, temporally-extended actions that determine the choices at each trial to each target. In particular, we can use the options framework [40] to define this hierarchical model. In this framework, an option *o* consist of three components: a policy *π*, a termination condition *β*,and an initiation set ℐ,

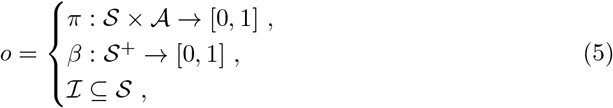

where 𝒮 is the state space, and 𝒜 is the primitive action space. If an option *o* = ⟨ℐ, *π, β*⟩ is taken, then actions are selected according to *π* until the option terminates stochastically according to *β*. This framework can be extended to semi-Markov options, where a history Ω over states, actions and rewards can be taken as input to the low level policy *π*. This is important because our foraging model will make use of this history.

In the foraging model, the primitive action space 𝒜 is the set of targets than can be chosen, the state space is a trivial one-state space (so initiation sets are trivial), and we define the option space as

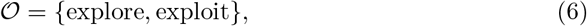

with each option defined the following way:

1. The exploit option has a lower level policy *π*_exploit_ that executes always the same action (given by the previous action from the history), and terminates with probability

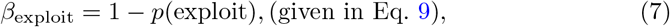

2. The explore option has a lower level policy *π*_explore_ that selects targets uniformly randomly, and terminates with probability

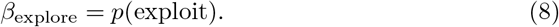

We define the termination conditions with a softmax rule, in a similar fashion to how the RL model randomly selects other targets, in the following way:

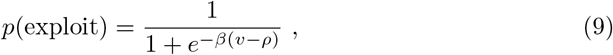

where *v* is a value that gets updated using Eq. 1 and the threshold *ρ* could be thought of as the expected value of any alternative (the background rate of reward in the environment [17]) or as a literal threshold for exploration. It is important to note that the foraging algorithm only tracks one value (which we can think of as the value of the exploit option). This way of interrupting or terminating options is a usual choice in the context of hierarchical actions and options [40, 61], where we decide to interrupt the execution of a temporally-extended action based on the value of that option compared to alternatives. With this space of options, we can now build a policy over options to define the foraging algorithm: whenever an option terminates, the other starts.

Because exploration ensured that a new option was chosen and value of exploiting it was unknown, the value of exploitation was reset to the threshold on each exploration trial.

### Extensions to Decision-Making Models

In addition to the standard formulations of the RL and foraging models introduced above, we considered a variety of extensions that are commonly used to improve the fit of RL models. First, we introduced an asymmetrical learning parameter, meaning a way for models to learn at different rates from rewards and omissions. Adding asymmetry often improves the fit of RL models [24, 62] and it improved the fit of both models here. In both the RL asymmetry and foraging asymmetry models, the delta-rule update was rewritten

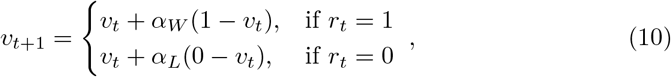

where *v* was the value of the chosen arm in the RL model or the value of exploitation in foraging. Both positive and negative learning rates were constrained between 0 and 1 in both models. Next, we introduced a decay parameter, meaning a way for some values to diminish over time without sampling. This approach is widely used in the reinforcement learning literature, where it has been interpreted as the indicative of limited memory resources. It consistently improves RL model fits [63–65] and improved the fit of both models here. In the reinforcement learning model, we implemented decay in the traditional way: instead of allowing the value of an unchosen option to carry forward unchanged in time, it decayed in proportion to delta (bounded at 0 and 1) as

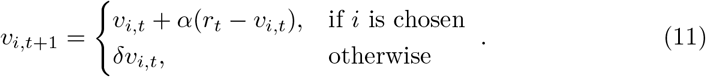

Although the foraging model has no unchosen values to decay, it does have a threshold parameter, *ρ*, which was previously modeled as static, but now allowed to change over time

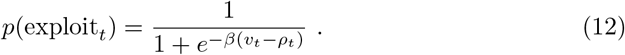

Specifically, *ρ* could now decay with each successive choice to the same option through updating it in a state-dependent way as a function of some decay parameter *δ*,

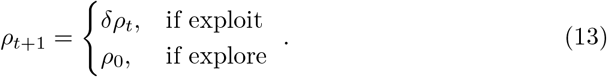

where *ρ*_0_ is the value at which *ρ* is initialized at time 0 or at the onset of exploration. Finally, we considered 2 models that included history dependence, meaning an outcome-independent tendency to repeat past behavior. Adding dependence on previous choices tends to improve the fit of RL models [4, 24], likely because of the strong tendency towards hysteresis in the choices of humans and other animals. Adding history dependence also improved the fit of both models here. In reinforcement learning, one common approach adds a choice kernel (*k*) for each option *i* which is updated in parallel with the value as

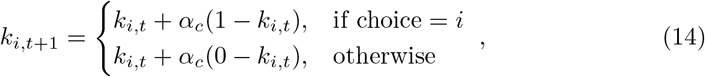

where *α*_*c*_ is the choice kernel learning rate parameter. The choice kernel value is then combined with value in the softmax decision rule,

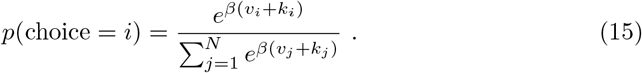

Thus far, we have added only 1 choice-history dependent parameter to this RL model, the *α*_*c*_ parameter, and this model is the history kernel 1 RL model referred to in the text. However, it is common to give choice history its own inverse temperature parameter *β*_*c*_. This is the history kernel 2 RL model,

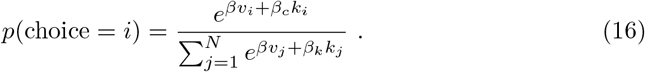

Although there was no natural way to add a choice history kernel to the foraging model, we could use a nearly identical approach to add a state history kernel to the foraging model. This approach adds hysteresis at the level of the exploitation state, rather than at the level of the choices themselves. To do this, we introduced a state history kernel that was updated alongside the value of exploitation as

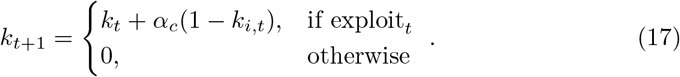

Meaning that the state history kernel was re-set to 0 when the animal explored. In the history-kernel 1 foraging model, this kernel was then added directly to the exponential in the decision rule,

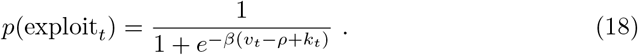

In the history-kernel 2 foraging model, the kernel was inserted, but scaled with its own inverse noise parameter, *β*_*k*_, as

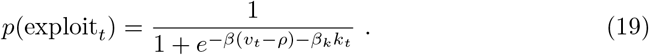

### Model Fitting Procedure

Maximum Likelihood Estimation (MLE) was performed to determine the parameter values of both standard models that most accurately described human behavioral data. The log likelihood (*LL*) was computed as follows:

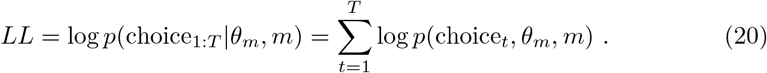

where *p*(choice_1:*T*_ |*θ*_*m*_, *m*) is the probability of all choices (from 1 to *T* trials) given the parameters *θ*_*m*_ of each model *m* with *θ*_*RL*_ = [*α, β*] and *θ*_*foraging*_ = [*α, β, ρ*] for each participant. To find the maximum likelihood parameters, the negative log-Likelihood was minimized using an optimization function (Experiment 1: Matlab fminsearch; Experiment 2-3: Python scipy.optimize.minimize). The optimization process was initiated multiple times with randomly selected starting parameters to avoid local minima (Experiment 1: 20 times, Experiment 2-3: 50 times). The set of parameters with the lowest negative Log-Likelihood were selected. The Akaike Information Criteria was calculated to identify the best fitting model, adjusting for differences in the number of parameters, and Akaike weights were calculated to estimate the relative ability of each model to minimize information loss.

### Simulations from Decision-Making Models

#### Performance Optimization

In order to determine if foraging had an advantage (or disadvantage) compared to RL in performance, we asked if the upper bound on performance was higher for foraging than it was for RL. This meant that we used simulation to identify the optimal parameter combination for each model in matched, randomly generated environments that matched the statistics of Experiment 1. We then compared the performance of these optimized models. Optimal parameters combination was selected so that it maximized the relative reward (minimizing the negative of the relative reward, Python scipy.optimize.minimize) while remaining within the participants’ parameters range.

#### Cohort simulation

When simulating data from the standard foraging and RL models, our goal was to produce datasets under each model that would resemble the set of human participants as closely as possible. To do this, we took the stochastic reward schedule experienced by each participant and simulated a new sequence of choices in that reward environment from the RL and foraging models. The simulations were generated using fitted parameters from the participant who experienced that environment.

### Exponential Mixture Model

We examined whether identifiable temporal patterns existed within the participants’ choice sequences. If a single time constant (probability of switching) governed the behavior, we would expect to see exponentially distributed inter-switch intervals. That is, the distribution of inter-switch intervals should be well described by the following model:

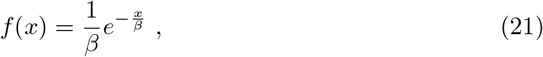

where *β* is the survival parameter of the model (mean inter-switch interval). Although the time between switch decisions was largely monotonically decreasing and concave upwards, the distribution was not well described by a single exponential distribution. Therefore, we next fit mixtures of varying numbers of exponential distributions (1-4) in order to infer the number of switching regimes in these choice processes. For continuous-time processes, these mixture distributions would be of the form:

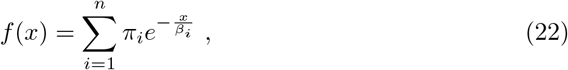

where 1 ≥ *π*_*i*_ ≥ 0 for all *π*_*i*_, and 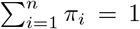. Here, each *β*_*i*_ reflects the survival parameter (average inter-switch interval) for each component distribution *i* and the *π*_*i*_ reflects the relative weight of each component. Because trials were discrete, we fit the discrete analog of this distribution: mixtures of 1-4 discrete exponential (geometric) distributions [66]. Mixtures were fit via the expectation-maximisation algorithm and we used standard model comparison techniques [67] to determine the most probable number of mixing components. Two regimes (log-likelihood = -31,048, 3 parameter) were significantly better fit to data than one (log-likelihood = -34,645, 1 parameter, likelihood ratio test: ratio = 7193.0, p *<* 10^*−*32^) and, while adding additional regimes continued to improve model fit (3 regimes: log-likehood = -30,893, 5 parameters; 4 regimes: log-likelihood = -30,884), improvement was already saturated at 2 regimes (Figure 5D).

### Hidden Markov Model

To determine when models and participants were exploring (versus exploiting), we used a hidden Markov model (HMM; [15, 24, 32, 36, 37]. In an HMM, choices (*y*) are ‘emissions’ that are generated by an unobserved decision process that is in some latent, hidden state (*z*). Latent states are defined by both the probability of making each choice (*k*, out of *N*_*k*_ possible options), and by the probability of transitioning from each state to every other state. Our model consisted of two types of states, an explore state and the exploit states. The emissions model for the explore state was the maximum-entropy distribution for a categorical variable, a uniform distribution:

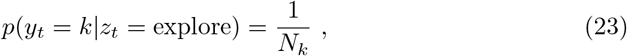

meaning that we made the fewest number of assumptions possible about the choices that were made during exploration in order to avoid biasing the model towards or away from any particular type of policy. However, modeling exploratory choices with a uniform distribution does not imply, require, or enforce random decision-making during these states [36, 37]. Because exploitation involves repeated sampling of each option, exploit states only permitted choice emissions that matched one option:

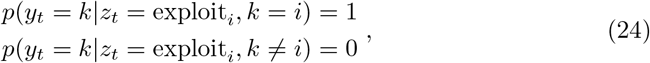

meaning latent states in this model are Markovian. The current state (*z*_*t*_) depends only on the most recent state (*z*_*t−*1_),

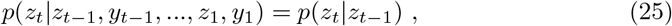

meaning that we can describe the entire pattern of dynamics in terms of a time-invariant transition matrix between 3 possible states (two exploit states and one explore state). This matrix is a system of stochastic equations that describe the one-time-step probability of transitioning between every combination of past and future states (*i, j*),

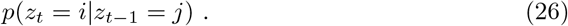

Because we had only a limited number of trials for each participant (300), parameters were tied across exploit states: each exploit state had the same probability of beginning (from exploring) and of sustaining itself. For conceptual reasons, the model also assumed that participants started in exploration and had to pass through exploration in order to start exploiting a new option, even if only for a single trial [15, 24, 32, 36, 37]. We have previously shown that models that lack these constraints by design tend to approximate them when fit to sufficiently large datasets [15, 24].

Because the emissions model was fixed, certain parameters were tied, the structure of the transmission matrix was constrained, and the initial state was specified, the final HMM had only two free parameters: one corresponding to the probability of exploring, given exploration on the last trial, and one corresponding to the probability of exploiting, given exploitation on the last trial. We have previously reported that this constrained model does not underperform an unconstrained model [15, 24]. and that unconstrained models tend to closely resemble to the constrained model when fit to large amounts of data [15].

The HMM was fit via expectation-maximization using the Baum Welch algorithm [68]. This algorithm finds a (possibly local) maxima of the complete-data likelihood. The algorithm was reinitialized with random seeds 20 times, and the model that maximized the observed (incomplete) data log likelihood across all the sessions for each animal was ultimately taken as the best. To decode latent states from choices, we used the Viterbi algorithm to discover the most probable a posteriori sequence of latent states.

### Explore/Exploit State Dynamics

In order to understand the dynamics of exploration and exploitation in the participants, RL models, and foraging models, we analysed the dynamics of the HMMs [24, 32, 37]. The fitted HMMs are a set of equations that describe probability of transitions between exploration and exploitation and vice versa, or the state dynamics of each agent. Methods from statistical thermodynamics can then be used to analyze these equations and generate insight into the potential energy of each state in each agent (Figure 4M).

In statistical thermodynamics, the potential energy of a state (*E*_*i*_) is related to the long-time scale probability of observing a process (here, a decision-maker) in that state (*p*_*i*_) via the Boltzman distribution,

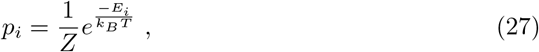

where *Z* is the partition function of the system, *kB* is the Boltzman constant, and *T* is the temperature. In a two-state system, the partition functions cancel out, the relative occupancy of the states is just a function of the difference in energy between them, and we can rearrange to express the difference in energy between two states as a function of the difference between them,

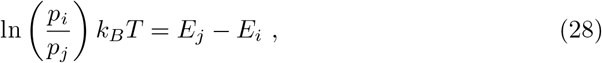

where *E*_*j*_ − *E*_*i*_ is the difference in the energetic depth between the two states (i.e. the Gibbs Free Energy), which is proportional to the log odds of each state, up to some multiplicative factor, *kBT*.

To calculate the probability of exploration and exploitation (*p*_*i*_ and *p*_*j*_), we solved for the stationary distribution *π*^*\**^ of the fitted HMMs, where *π*^*\**^ is the probability distribution that satisfies

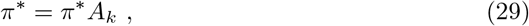

where *A*_*k*_ is the transition matrix for agent *k*. If it exists, this distribution is a (normalised) left eigenvector of *A*_*k*_ with an eigenvalue of 1, so we solved for this eigenvector to determine the stationary distribution over explore and exploit states for each agent. We then took an average of these stationary distributions across all sessions for each species, and plugged these back into the Boltzman equations to calculate the relative energy (depth) of exploration and exploitation in (Figure 4M).

In order to calculate the height of the energetic barrier between exploration and exploitation, we built on the Arrhenius equation from chemical kinetics that relates the rate of transitions (k) between some pair of states to the activation energy required to affect these transitions (Ea):

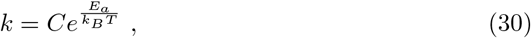

where *C* is a constant pre-exponential factor and *kBT* is again the product of temperature and the Boltzman constant. Rearranging to solve for activation energy yields our equation for the height of the energy barrier,

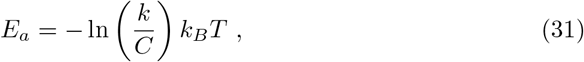

which has an obvious similarity to the Boltzman distribution illustrated earlier, where the relative depth of each state was proportional to the probability of occupying each state and the activation energy is now proportional to the rate of transitions between states.

To create the dynamical landscape graphs (Figure 4M), transition matrices were calculated individually for all participants and for simulated data from both foraging and RL algorithms. Energy measurements were then averaged within each class of agents. Note that our approach only identifies the energy of three discrete states (an explore state, an exploit state, and the peak of the barrier between them). These are illustrated by tracing a continuous potential through these three points to provide a physical intuition for the differences in explore/exploit dynamics between models and participants.

### Foraging Index

In order to determine if individual participants were more foraging-like (versus RL-like) in their approach to the task, we calculated the foraging index as the difference in the individual model likelihood (L) between the best-fitting foraging model (F*) and the best-fitting RL model (RL*) for each participant i:

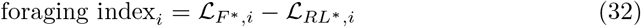

## Resource Availability

## Material Availability

This study did not involve generating new material.

## Data and Code Availability

Requests for Data and Code should be directed to and will be fulfilled by Lead Contacts. Data and code will be made publicly available upon acceptance of this article.

## Author contributions

The author contribution matrix was adapted from [69].

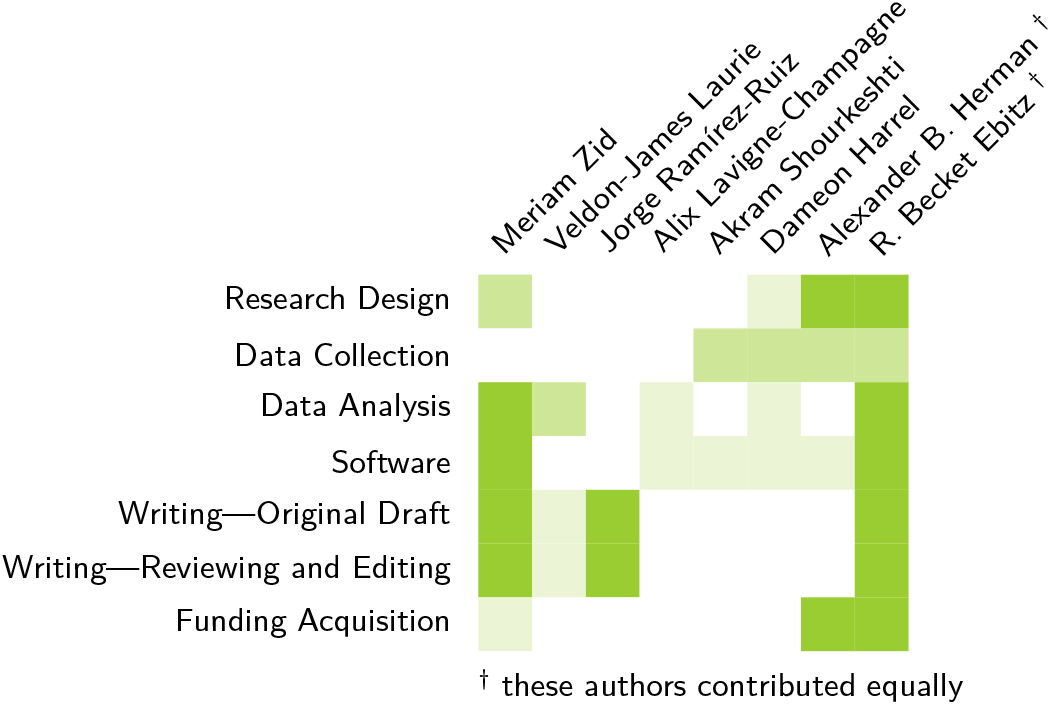

## Acknowledgements

The authors would like to thank Ossama Ghenissa, Rishabh Singhal, and Nicola Grissom for insightful discussion, Devin H. Keheo for feedback on an earlier draft of the manuscript, and Cathy Chen and Colin Meyer for invaluable technical support. This project was supported by an NSERC Discovery Grant (RGPIN-2020-05577; RBE), the Research Corporation for Science Advancement & Frederick Gardner Cottrell Foundation (Project #29087; RBE), the Canada Research Chair Dynamics of Cognition (FD507106; RBE), a Research Fellowship from the Jacobs Foundation (RBE), the Centre Interdisciplinaire de Recherche sur le Cerveau et l’Apprentissage (CIRCA; M.Z.), and the National Institutes of Health (R21 MH127607 and K23 MH050909; A.B.H.).

